# The bacterial DNA binding protein MatP involved in linking the nucleoid terminal domain to the divisome at midcell interacts with lipid membranes

**DOI:** 10.1101/428714

**Authors:** Begoña Monterroso, Silvia Zorrilla, Marta Sobrinos-Sanguino, Miguel Ángel Robles-Ramos, Carlos Alfonso, Bill Söderström, Nils Y Meiresonne, Jolanda Verheul, Tanneke den Blaauwen, Germán Rivas

## Abstract

Division ring formation at midcell is controlled by various mechanisms in *Escherichia coli*, one of them being the linkage between the chromosomal Ter macrodomain and the Z-ring mediated by MatP, a DNA binding protein that organizes this macrodomain and contributes to the prevention of premature chromosome segregation. Here we show that, during cell division, just before splitting the daughter cells, MatP seems to localize close to the cytoplasmic membrane, suggesting that this protein might interact with lipids. To test this hypothesis, we investigated MatP interaction with lipids *in vitro*. We found that MatP, when encapsulated inside microdroplets generated by microfluidics and giant vesicles, accumulates at phospholipid bilayers and monolayers matching the lipid composition in the *E. coli* inner membrane. MatP binding to lipids was independently confirmed using lipid coated microbeads and bio-layer interferometry assays. Interaction of MatP with the lipid membranes also occurs in the presence of the DNA sequences specifically targeted by the protein but there is no evidence of ternary membrane/protein/DNA complexes. We propose that the interaction of MatP with lipids may modulate its spatiotemporal localization and its recognition of other ligands.

**IMPORTANCE:** The division of an *E. coli* cell into two daughter cells with equal genomic information and similar size requires duplication and segregation of the chromosome and subsequent scission of the envelope by a protein ring, the Z-ring. MatP is a DNA binding protein that contributes both to the positioning of the Z-ring at midcell and the temporal control of nucleoid segregation. Our integrated *in vivo* and *in vitro* analysis provides evidence that MatP can interact with lipid membranes comprising the phospholipid mixture in the *E. coli* inner membrane, without concomitant recruitment of the short DNA sequences specifically targeted by MatP. This observation strongly suggests that the membrane may play a role in the regulation of the function and localization of MatP, which could be relevant for the coordination of the two fundamental processes in which this protein participates, nucleoid segregation and cell division.

## INTRODUCTION

Bacterial division is achieved through the assembly of a protein machinery into a membrane anchored ring that splits the cell generating two daughter cells with equal genomic information (1). The scaffold for the involved proteins is the self-assembling protein FtsZ. The need of a precise localization of this Z-ring in the middle of the cell is fulfilled by different mechanisms evolved in bacteria, the canonical ones being the Min system and nucleoid occlusion (2). An additional mechanism contributing to Z-ring positioning is the linkage between the Ter macrodomain of the chromosome and the Z-ring (Ter linkage (3)). While the two first systems exert their action through blockage of productive FtsZ assembly at certain locations, namely the vicinity of the nucleoid and the cell poles, the last one is a positive mechanism promoting assembly of the division machinery nearby the replication terminus region of the chromosome (4).

The Ter linkage consists of three proteins, MatP, ZapB, and ZapA which form a complex that links the chromosome to the Z-ring. MatP, a DNA binding protein, was identified by Mercier and coworkers (5), who showed that it is the main organizer of the Ter macrodomain of the chromosome, preventing its premature segregation through specific interaction with a short palindromic DNA sequence (*matS*) repeated 23 times within this macrodomain. There are no *matS* sequences outside the Ter macrodomain, which is in turn devoid of the sequences targeted by SlmA, the other DNA binding protein avoiding, through nucleoid occlusion, aberrant Z-ring positioning (6). It was recently found that, upon binding to the *matS* sites, MatP displaces MukBEF from the Ter domain (7) promoting the formation of a unique chromosomal region. The Ter domain progressively shifts towards the cell centre along the cell cycle (8) and by binding to ZapB remains localized at midcell during division in slowly growing cells (9, 10). During the last cell division stage the Ter macrodomain is segregated into each daughter cell while they separate. The cell division protein FtsK forms probably at this stage a hexameric DNA translocase that moves about 400 bp towards the *dif* sites close to the terminus of the chromosome while displacing MatP from its *matS* sites (11, 12) to assist in the segregation of the termini. The molecular mechanisms by which this last step of chromosome segregation and daughter cell separation are coordinated remain largely unknown.

In this work, we observed that MatP moves away from the division site near the end of the cell division cycle, leaving a still intact divisome including ZapB at midcell. Indeed, also the colocalization with the nucleoids seemed to be at least partly lost and MatP was often observed close to the cytoplasmic membrane. On the basis of these findings, we postulated that MatP could bind to the lipids in the cytoplasmic membrane. To verify this hypothesis, we investigated the interaction of MatP with lipids *in vitro*. We reconstructed the purified protein MatP inside microfluidics microdroplets and giant unilamellar vesicles (GUVs), and found a significant preference of the protein for the lipid membrane compared with the lumen of the container. The shift of the protein towards the membrane occurred also in the presence of an oligonucleotide containing the DNA sequence targeted by MatP, *matS*, but no accumulation of this sequence was detected at the membrane. Parallel experiments based on complementary biochemical approaches further supported the interaction of MatP with lipids. Our results indicate that MatP constitutes another example of a protein involved in division able of recognizing both nucleic acid sequences and lipid membranes, as previously described for MinD (13) and *Bacillus subtilis* Noc (14). Furthermore, we propose that the membrane binding of MatP serves to free the *matS* sites close to the *dif* site that is needed by FtsK to help the segregation of the termini into the two daughter cells.

## RESULTS

### MatP localizes in between the nucleoid and ZapB at the end of the cell division cycle

To investigate what the exact sequence of events is during the process of cell division, we previously analysed the localization of a large number of cell division proteins in steady state slowly growing cells (15, 16). The advantage of slowly growing cells is that they do not have multiple replication forks at least during the major part of their division cycle. When *E. coli* cells are grown to steady state, their length correlates well with the cell division cycle age. We have now investigated, as part of the proteins that are involved in the coupling of cell division and chromosome segregation, the localization of the nucleoids in relation to that of MatP and the protein complex responsible for division (divisome). MG1655 cells expressing MatP-mCherry (17) from the original locus in the chromosome were grown in minimal medium to steady state. In these cells the localization of its divisome partner ZapB and the divisome protein FtsN that marks the presence of a complete division machinery were determined by immunolabelling.

To be able to dissect what happens to the localization of these three proteins and the nucleoid (stained by DAPI), more than 16000 cells were imaged and analysed. MatP-mCh and ZapB colocalize during most of the cell division cycle and FtsN arrives later at midcell (**Fig. 1A**) as described (8, 15, 17). The concentration of MatP is constant during the cell cycle (**Fig. S1A**). The number of MatP dimers, species assumed based on previous structural *in vitro* data (17), per average cell in minimal medium was determined to be 180 (18). Using this number and the determined extra fluorescence at midcell (FCPlus (16)), the number of MatP dimers was calculated to be 60 in the foci at midcell at 80% of the cell division cycle age (**Fig. S1B**). MatP localizes in young cells as a diffuse focus, which moves toward the cells centre during the cell division cycle where it forms a more distinct concentrated focus (**Fig. 1AB** and **Fig. S1C**). When determining the position of the brightest pixel in the MatP foci, they seem to localize consistently close to the length axis of the cell (**Fig. S1D**) as was reported (19). However, after 90% of the cell division cycle MatP-mCh moves away from the divisome, whereas ZapB and FtsN remain almost till the cells are completely divided (**Fig. 1A**). Interestingly, inspection of the deeply constricting cells suggested that MatP is not following the nucleoid that is segregating but remains in between ZapB and the nucleoid. This suggests that at least part of the MatP protein is not binding to the Ter domain any longer and also not to ZapB.

**Fig. 1.**
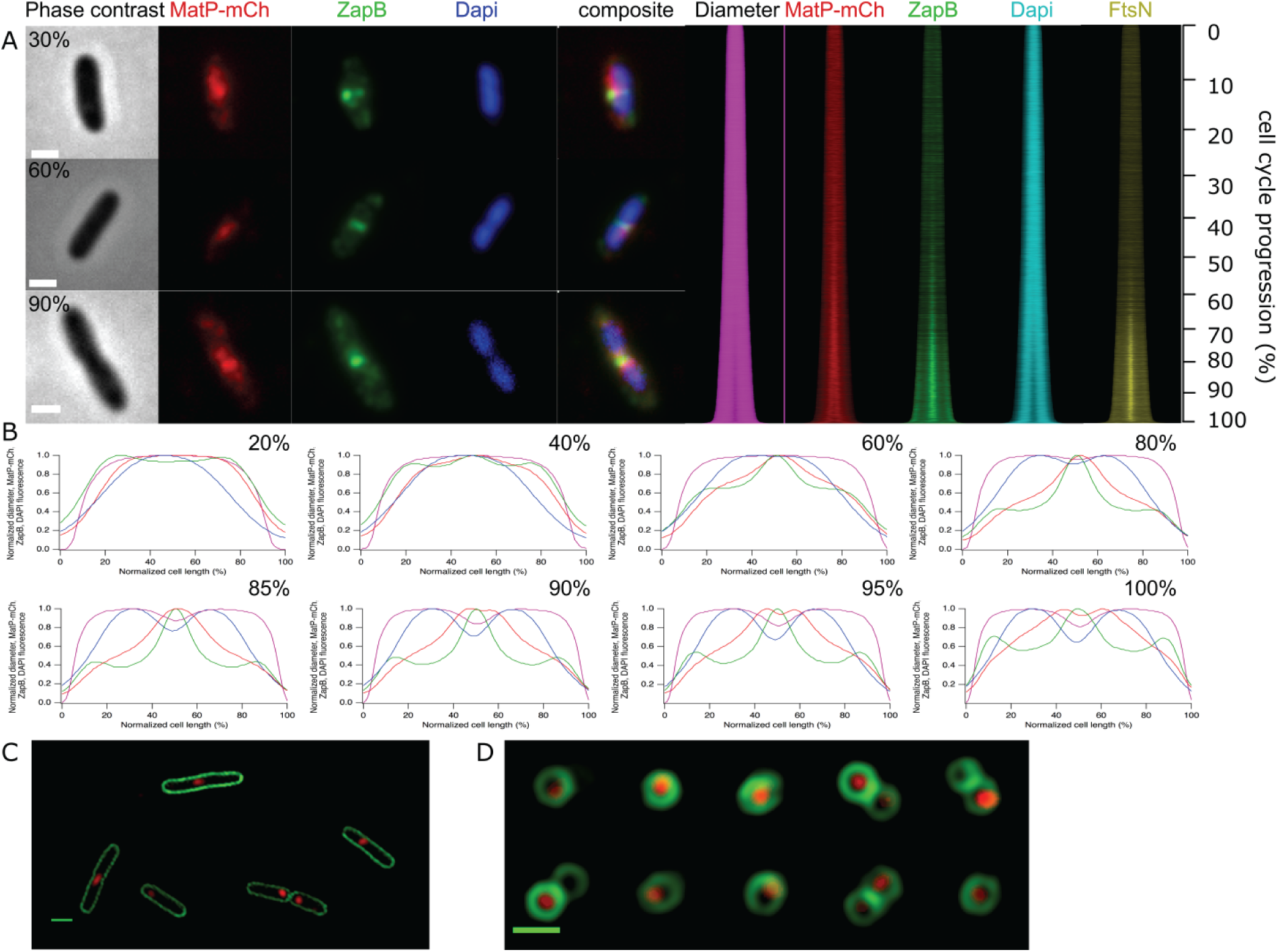
Localization of MatP as a function of the cell division cycle. (A) A representative example of a cell of age class 30%, 60% and 90% from top to bottom and from left to right: Phase contrast and fluorescence images of MatP-mCh, immunolabelled ZapB and DAPI stained nucleoids are shown. The map of profiles shows in the same order the diameter determined in the phase contrast images, and the fluorescence as a function of cell length. The numbers on the right show the relation between length and cell division cycle age. (B) Peak normalized average profiles from the maps of the diameter and fluorescence were plotted against the normalized cell length in age bins of 0–20%, 20–40%, 40–60%, 60–80%, 80–85%, 85–90%, 90–95% and 95–100%. The age class with the smallest number of cells, *i.e*. 95–100% still contains 592 cells. In total 16796 cells were analysed. (C) SIM images of life MG1655 *matP-mCh::kan* transformed with the plasmid pXL28 that expresses the integral membrane protein fusion mNeonGreen-(GGS)_2_-GlpT. The cells had been grown in Gb4 minimal medium at 28°C and induced for 2 mass doublings with 30 μM IPTG. (D) SIM images of upright positioned cells grown as in C. The 10 cells shown are all from a single image without selection and grouped together to reduce the figure size. All scalebars equal 2 μm.

Since we observed the signal of MatP often close to the membrane of the new poles, we wondered whether MatP might bind lipids, like was observed for other proteins binding to the chromosome such as the Noc protein (14). To determine whether MatP-mCh colocalized with the cytoplasmic membrane, we transformed MG1655::MatP-mCh with a plasmid pXL28 that expresses the integral membrane protein fusion mNeonGreen-(GGS)_2_-GlpT (20). Cells were grown to steady state and the colocalization of MatP and GlpT was determined by the colocalization of the fluorescence of both proteins using the Pearson coefficient (21) as a function of the cell division cycle (**Fig. S2A**). The same strain without plasmid was used to determine the amount of overlap with the mCh channel due to autofluorescence. The Pearson coefficient did increase from 0.18 ± 0.13 in cells without the membrane staining mNG-GlpT to 0.32 ± 0.13 in cells that did express the protein, indicating some overlap. Because MatP-mCh consisted of one or two foci per cell and the mNG-GlpT was distributed evenly in the cell membrane not a large overlap was to be expected and no striking difference in the very old cells was observed (**Fig. S2A**).

An increase in the Pearson coefficient due to binding of single MatP-mCh molecules (*i.e*. not bound to DNA) to lipids cannot be discarded, since they are not observable by wide field fluorescence microscopy. Trying to discriminate between these options we used structured illumination microscopy (SIM) of cells (**Fig. 1C**) immobilized in an upright position (**Fig. 1D**) using an agar-pad with a range of micrometre-sized holes and looked at the colocalization of MatP and GlpT. A collection of cells taken from one image (no selection) is shown in **Fig. 1D**. Many foci localized in the middle of the circumference of the cell short axis and some colocalization of MatP and the membrane was observed, reinforcing the idea that MatP could interact with lipids. The resolution of the microscope and the intensity of the mCherry signal were not sufficient to discriminate binding of single mCherry molecules to the membrane. Therefore, we decided to investigate the membrane binding of MatP further *in vitro*.

### MatP accumulates at the lipid boundaries of microdroplets and vesicles

With the aim to investigate whether MatP had lipid affinity we encapsulated the protein, using microfluidics based technology, inside microdroplets as cell mimic systems surrounded by a lipid boundary resembling that of the *E. coli* inner membrane. MatP (with a tracer amount of MatP-Alexa 488) was included in one of the aqueous streams, the other one being buffer (**Fig. 2A**). Microdroplets were formed when the aqueous solutions met the continuous phase, constituted by the *E. coli* lipids dispersed in mineral oil, at the production junction of the microchip. Interestingly, according to the confocal microscopy images of the samples, MatP was mostly located at the lipid interface inside the microdroplets, as reflected by the intensity profiles (**Fig. 2A**). Interaction of the protein with lipids was also found when the solution encapsulated inside microdroplets contained crowding agents like Ficoll or dextran together with MatP (**Fig. S3AB**). No preference for the lipid boundary was observed when the free Alexa 488 dye was encapsulated (**Fig. S3C**).

**Fig. 2.**
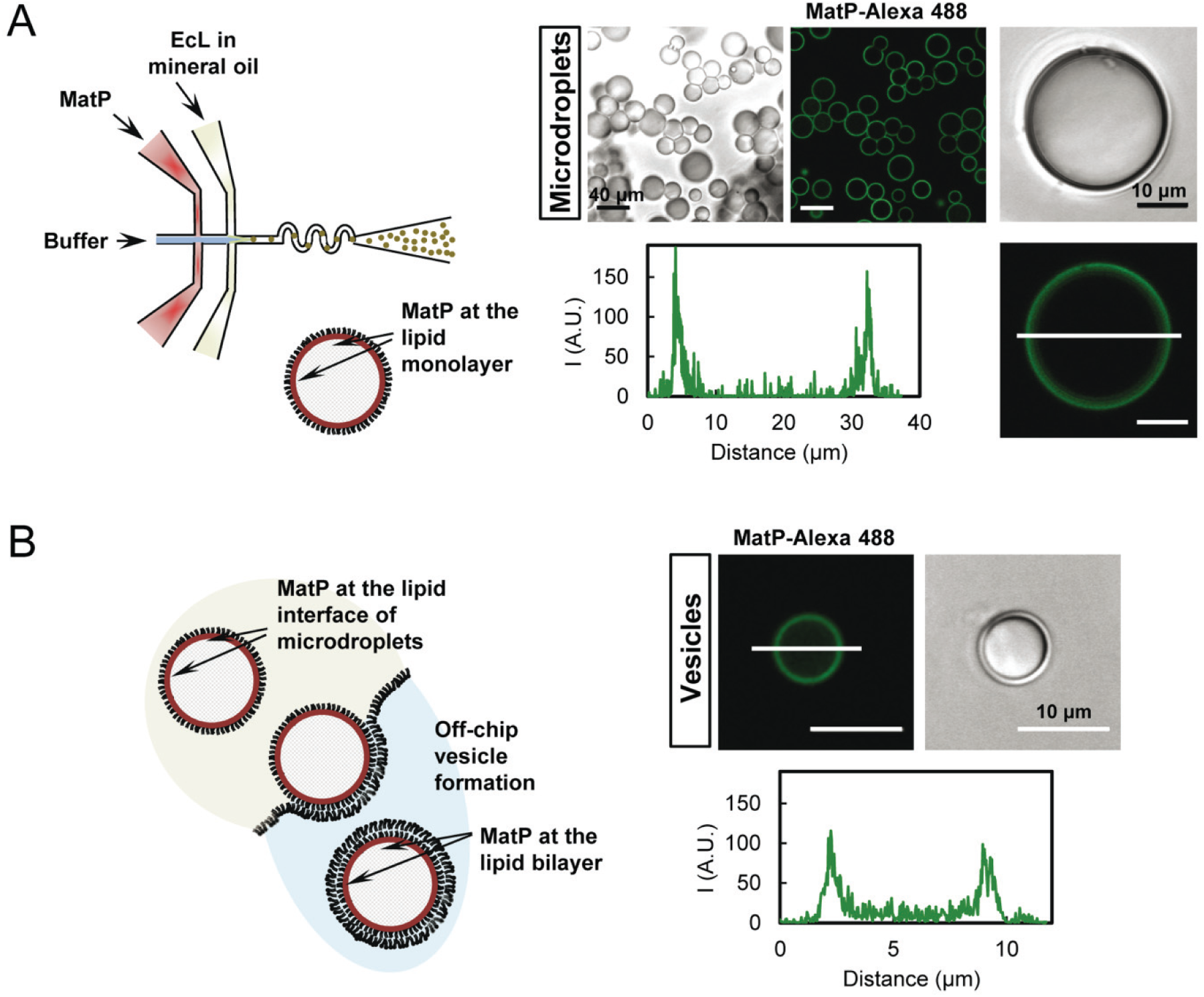
Microfluidic encapsulation of MatP inside microdroplets stabilized by the *E. coli* lipid mixture and GUVs formed from them. (A) Scheme of the encapsulation setup and of the distribution of species within the droplet (left). Representative confocal and transmitted images of the microdroplets containing MatP (3.5 μM), and intensity profile corresponding to the green channel (MatP-Alexa 488, 1 μM), obtained across the line as drawn in the image (right). (B) Illustration of the step determining vesicle formation from the droplets with MatP and of the distribution of species within the GUVs (left). Representative confocal and transmitted images of GUVs and intensity profile corresponding to the green channel (MatP-Alexa 488), obtained across the line as drawn in the image (right). Vesicles contained 150 g·L^−1^ Ficoll.

After this observation, we wanted to study whether MatP interaction with the lipids also occurred when the lipid boundary was a bilayer, which provides a better cell-like system, instead of the monolayer surrounding the microdroplets. For this purpose, the microdroplets obtained by microfluidics were converted into GUVs, using a procedure based on the droplet transfer method (22), as previously described (23). The droplets acquired the bilayer upon transition from an oil phase to an aqueous solution through an interface coated with oriented lipids (**Fig. 2B**). The crowding agent Ficoll was encapsulated alongside with MatP and the osmolarity of the solutions was adjusted to improve vesicle integrity and yield. Confocal images of the samples and the corresponding intensity profiles showed a remarkable shift of green labelled MatP towards the lipid membrane of the GUVs (**Fig. 2B**). Binding to lipids also occurred when MatP was externally added to GUVs (**Fig. S4**).

These results showed that the division protein MatP interacts with lipid monolayers or bilayers resembling the composition of the *E. coli* inner membrane when encapsulated inside micron sized cytomimetic containers.

### MatP interacts with *E. coli* lipid bilayers at submicromolar concentrations

To quantify the interaction of MatP with lipid membranes, bio-layer interferometry assays were conducted using biosensor tips coated with the *E. coli* lipid mixture. Addition of the protein resulted in a shift in the incident light directed through the biosensor, indicative of binding (**Fig. 3A**). A dose-response curve obtained by varying the concentration of MatP showed that, above 10 nM, the biosensor signal associated with binding increases with protein concentration, being saturated at around 1 μM MatP (**Fig. 3B**). Analysis of this curve with an empirical Langmuir adsorption equation, with no assumption about the mechanism or stoichiometry of the binding, rendered a *c*_50_ value of 97 nM with a well-defined upper limit (**Fig. S5**), corresponding to the concentration of MatP at which half of the maximum response signal was observed.

**Fig. 3.**
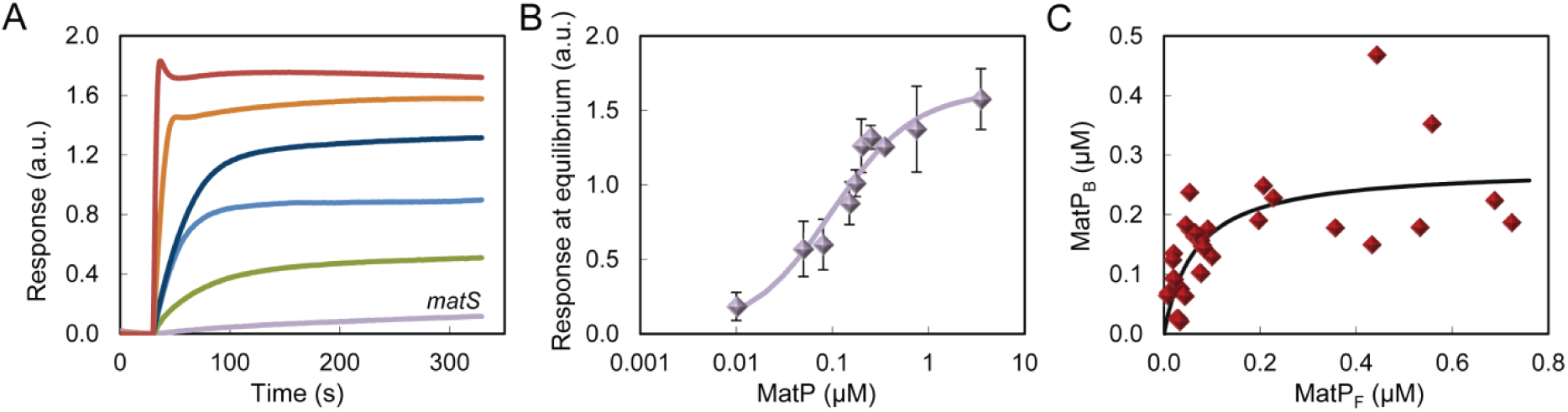
Binding of MatP to *E. coli* lipids by bio-layer interferometry or using lipid coated microbeads. (A) Representative profiles of the binding of MatP at increasing concentrations obtained by bio-layer interferometry. From bottom to top, 100, 150, 250, 750 nM and 3.5 μM. The profile obtained for *matS* is shown for comparison. (B) Dose-response curve obtained as a function of the concentration of MatP. Solid line is the best fit according to the model explained in the main text rendering the following parameter values: *c*_50_ = 97 nM and *y_max_* = 1.6. (C) MatP binding to *E. coli* lipid coated microbeads plotted as a function of the concentration of free MatP. Symbols are the data, and the solid line is the best fit according to the model explained in the main text rendering the parameter values *c*_50_ = 65 nM and *y_max_* = 280 nM. [Beads] = 35 g·L^−1^ (62 μM accessible lipid). MatP was labelled with Alexa 488.

The binding of MatP to lipids was also ascertained through co-sedimentation assays using microbeads coated with the *E. coli* lipids mixture and MatP labelled with Alexa 488. Significant depletion of the protein was observed after incubation with the beads and centrifugation, and the amount of protein bound increased with the concentration of protein at constant concentration of lipids (**Fig. 3C, Fig. S6**). Observation of the microbeads after incubation with the green labelled protein by confocal microscopy confirmed the interaction (**Fig. S6**). As in the bio-layer interferometry experiments, the binding isotherm obtained by plotting the concentration of protein bound to the beads against the concentration of free protein was analysed using the empirical Langmuir isotherm (**Fig. 3C**). This analysis rendered a *c*_50_ of 65 nM, again with a well-defined upper limit (**Fig. S6**), close to the midpoint of the response curve obtained by bio-layer interferometry.

The bio-layer interferometry assays and the lipid coated microbead experiments further supported the interaction of MatP with lipids, showing that it occurs at submicromolar concentrations of the protein.

### MatP does not recruit *matS* to the membrane

As MatP is a DNA binding protein, we asked if it was still able of binding to the *E. coli* lipids in the presence of oligonucleotides containing its specific binding sequence, *matS*. To approach this question, we first characterized the protein/DNA complexes in the working buffer used to encapsulate MatP. MatP behaved as a dimer, as determined by sedimentation and light scattering (see analysis of MatP·*matS* complexes under Supplementary information and **Fig. S7B**), in good agreement with previous data (17). The stoichiometry of the MatP *matS* complex was determined to be two monomers of MatP and one molecule of the *matS19* target (**Fig. S7**), again in agreement with crystallography analysis (17). The *Kd* for the interaction, determined by fluorescence anisotropy under our experimental conditions, was 15 ± 2 nM, in dimer units (see analysis of MatP·*matS* complexes under Supplementary information and **Fig S7**).

We next encapsulated MatP alongside with m*atS*19 inside microdroplets, to analyse the influence of the oligonucleotide on MatP interaction with the lipids. Encapsulation of MatP (with a tracer amount of MatP-Alexa 488) and *matS*-Alexa 647 showed that location of MatP, almost exclusively at the lipid boundary of the microdroplets or GUVs, was not altered by the presence of *matS*, while the DNA, in turn, remained homogeneously distributed in their lumen (**Fig. 4AB**). Remarkably, the intensity profiles showed a drop of the red signal at the edges of the vesicle where the green signal corresponding to MatP reaches its maximum. This strongly suggests that MatP at the membrane is not bound to the DNA. The concentrations of MatP and *matS19* in these experiments were well above their *Kd* of interaction and we used a protein (monomer) molar excess relative to the DNA concentration above 2-fold, to ensure formation of the 2:1 complex previously characterized in solution (see above). The same results were found either by including MatP and *matS* in two independent streams, triggering complex formation shortly before encapsulation, or by encapsulating the preformed complex (*i.e*. MatP and *matS* together in the two streams). Additional experiments in which the fluorescein labelled *matS* used in the fluorescence anisotropy binding titrations and unlabelled MatP were encapsulated showed, again, that the DNA remained in the lumen of the microdroplets (**Fig. S8A**). The images obtained in this case were indistinguishable from those corresponding to the encapsulation of fluorescein labelled *matS* alone (**Fig. S8B**). These experiments evidenced that MatP still binds to the lipid monolayers or bilayers of cell-like containers in the presence of *matS*, although there was no sign of concomitant DNA recruitment to the lipid edge.

**Fig. 4.**
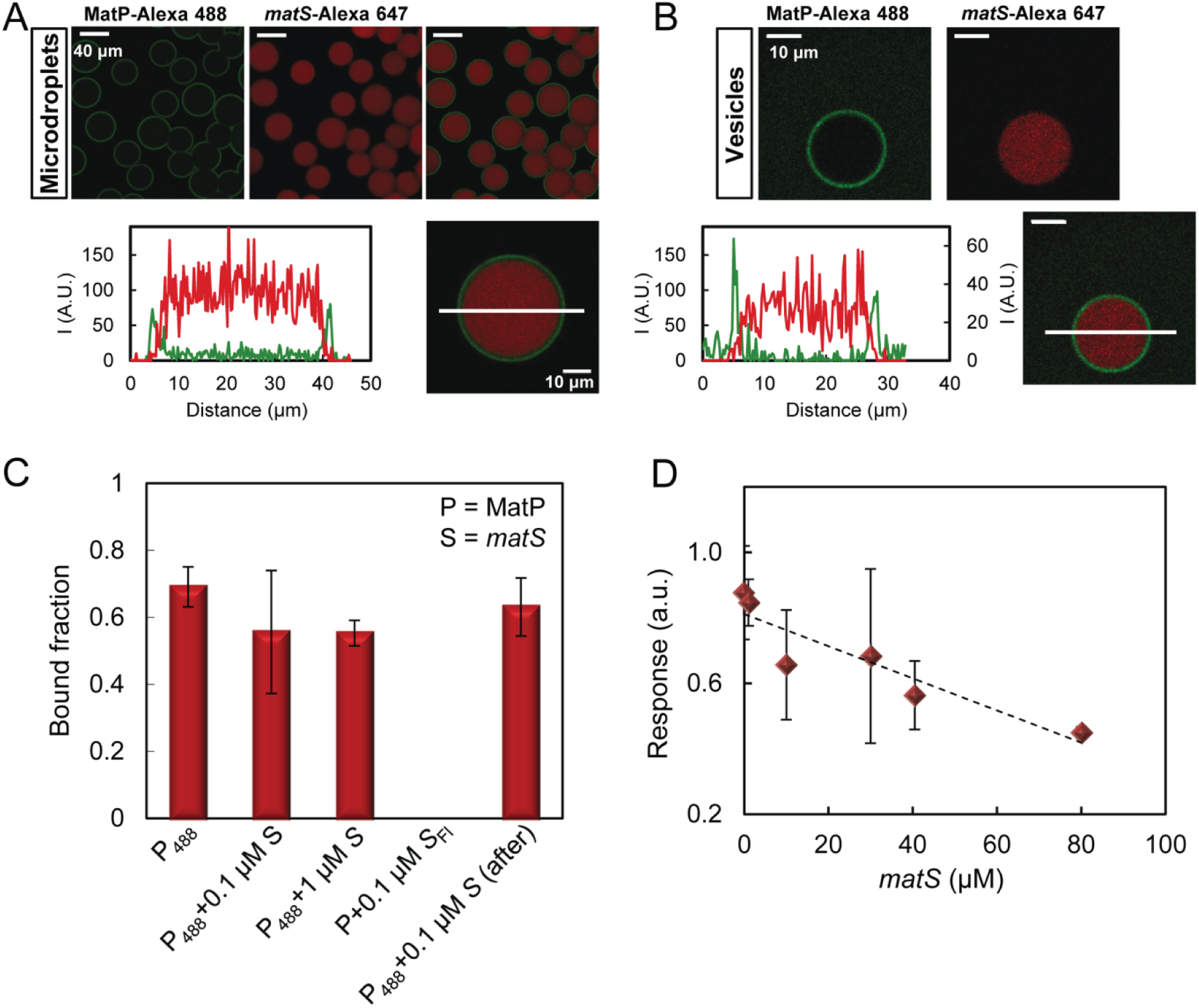
Binding of MatP to lipids in the presence of *matS*. (A, B) Representative confocal images of microdroplets and GUVs, respectively, stabilized by the *E. coli* lipids mixture containing MatP and *matS*, and intensity profiles below. Vesicles also contained 150 g·L^−1^ Ficoll. Profiles correspond to the green (MatP-Alexa 488, 1 μM) and red (*matS*-Alexa 647, 1 μM) channels, obtained across the line as drawn in the images. The concentrations of MatP and *matS* were 3.5 and 1.4 μM respectively. (C) Effect of *matS* on MatP binding to lipid coated beads (30 g·L^−1^, 53 μM accessible lipid). MatP concentration was 0.25 μM. *P*, *P_488_, S* and *S_Fl_* stand for MatP, MatP labelled with Alexa 488, *matS* and *matS* labelled with fluorescein, respectively. For the measurement corresponding to the bar on the far right, *matS* was added to MatP already bound to the lipid. (D) Competition of *matS* with the lipids for binding to MatP as observed by bio-layer interferometry. The concentration of MatP was 0.15 μM.

Next, we probed the influence of *matS* on the binding of MatP to lipids using microbeads coated with the *E. coli* lipid mixture and through bio-layer interferometry. Addition of 0.1–1 μM unlabelled *matS* prior or after incubation of MatP with the microbeads did not significantly modify the fraction of MatP-Alexa 488 bound with respect to that in the absence of *matS* (**Fig. 4C**). Parallel experiments using fluorescein labelled *matS* and unlabelled MatP showed that the DNA did not bind to the lipids alongside with MatP, in good agreement with the images of the complex encapsulated inside lipid vesicles or microdroplets (**Fig. 4C**).

Bio-layer interferometry assays conducted to measure the binding of MatP in the presence of constant 1 μM concentration of *matS* rendered isotherms of binding superimposable, within error, with those obtained in the absence of *matS* (**Fig. S5**, *c*_50_ = 76 nM), and no significant interaction with the lipids was detected for *matS* alone (**Fig. 3A**). The signal of binding of MatP (150 nM) to the lipids remained relatively insensitive to the concentration of *matS* below 30 μM, showing a decrease at higher concentrations (**Fig. 4D**) that suggests competition between the lipids and the DNA for binding to the protein. These experiments show that the interactions of MatP are relatively insensitive to the presence *matS*, which only seems to compete with lipid binding at high concentrations.

## DISCUSSION

Here we have found that the protein of the Ter linkage MatP interacts with membranes matching the lipid composition of the *E. coli* inner membrane, as shown by encapsulation in cell-like containers, co-sedimentation with lipid coated microbeads and bio-layer interferometry assays. Although MatP presents dual recognition of lipids and nucleic acid sequences, we have not found any indication supporting the formation of ternary complexes, strongly suggesting that both types of ligands may be mutually exclusive, which is also illustrated by the predominantly axial localization of the MatP foci. Competition between lipids and DNA for the same region of the protein, or lipid-induced changes in the association state and/or conformation of MatP hampering DNA binding, may explain the lack of DNA recruitment to the membrane.

Recent studies have revealed that, like MatP, other proteins binding to the bacterial chromosome are also able of interacting with lipid membranes. Examples of these proteins are the nucleoid occlusion Noc from *Bacillus subtilis* (14), a negative modulator of Z-ring assembly, and SeqA from *E. coli*, a protein involved in the sequestration of replication origins (24). Along the same line, the nucleoprotein complexes of SlmA, the factor counteracting Z-ring formation around the chromosome in *E. coli*, seem to be brought close to the membrane (4, 25, 26), possibly through transertional linkages (25) and/or biomolecular condensation (27). Conversely, well-known membrane associated proteins like MinD from the Min system (28) have also been shown to interact, in a non-sequence specific manner, with chromosomal DNA (13).

Membrane binding of MatP may serve to sequester the protein from the chromosome under conditions in which its positive regulation of Z-ring formation is no longer required and even might obstruct the function of proteins like FtsK. FtsK is needed for the deconcatenation of sister chromosomes and helps to segregate the termini into each daughter cell (9, 29). FtsK, part of the divisome (30), is one of the fastest DNA translocases (31). Membrane binding of MatP released by FtsK during this relatively short time interval might function to prevent rebinding to the *mat*S sites close to the *dif* site. We propose that *matS* and lipid competition for MatP assist in the segregation of the *dif* region by FtsK during the last step of septum closure (**Fig. 5**). The subsequent release from the membrane to bind again the *mat*S sites might be assisted by the oscillation behaviour of MinD that displaces proteins from the membrane surface of the new poles (32, 33). It is well known that the Min system oscillates between the old poles and the newly formed septum before the daughter cells have separated (34).

**Fig. 5.**
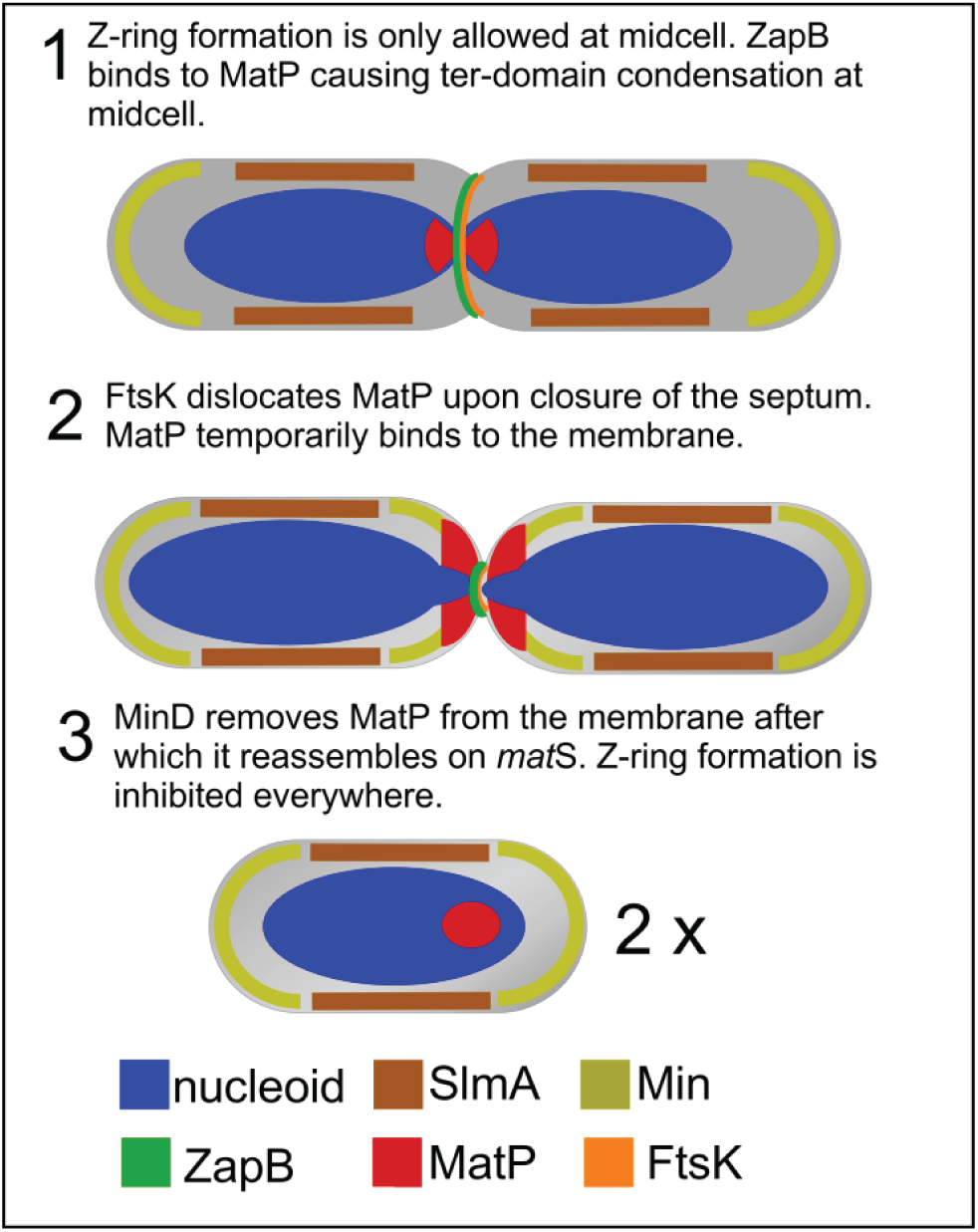
Hypothetical model for the MatP dissociation from the *matS* sites and its binding to the cytoplasmic membrane. 1. ZapB is binding MatP at midcell causing the 23 *matS* sites to cluster together and ensures that the terminus remains at midcell. Z-ring formation is inhibited at the old poles by the Min system and in the cylindrical part of the cell close to the bulk of the nucleoid, but not in the Ter-domain, by the nucleoid occlusion protein SlmA. 2. The nucleoids are segregating and MatP is pulled away from ZapB. At the same time the terminus is bound by FtsK that displaces MatP from *matS* sites by translocation of the DNA near the terminus, which allows final segregation of the nucleoids into the daughter cells. 3. MinD removes MatP from the membrane after which it reassembles on *matS*. Z-ring formation is inhibited everywhere by the Min system and the nucleoid occlusion protein SlmA.

The main difference between the dual recognition of lipids and DNA by MatP and the other site selection proteins (Noc and MinD) is that the latter can simultaneously bind DNA and lipids. Furthermore, in the particular case of Noc, binding to DNA activates in turn the subsequent interaction with the membrane (14). In bacillary bacteria, DNA-membrane interactions may aid in the localization of the bacterial cell centre, where the strength of these interactions decreases, and Z-ring assembly is favoured (35). By bridging the chromosome and the membrane, negative regulators of FtsZ polymerization such as Noc would exclude FtsZ from those areas biasing FtsZ assembly to the midcell (14). In contrast, physical connection of the chromosome with the membrane through MatP may interfere with its positive regulation of FtsZ assembly that contributes to division ring positioning. In the last step of binary fission that requires deconcatenation of sister chromosomes and closure of the septum, MatP’s presence might not be beneficial any longer. Therefore, it is displaced to the membrane to prevent immediate rebinding to the Ter domain, which would happen otherwise given its high affinity for these sites.

Since its identification, the function of MatP and its modulation in the context of division have been traditionally linked to its specific binding to DNA sequences within the Ter macrodomain or to its interaction with other proteins such as ZapB. Our findings strongly suggest that, in addition to protein-nucleic acid and protein-protein interactions, protein-lipid recognition should also be taken into account in the analysis of the function of MatP. Further work will be required to elucidate the precise mechanisms of these protein-membrane interactions and the factors influencing them.

## METHODS

### Chemicals and reagents

Polar extract of *E. coli* phospholipids, from Avanti Polar Lipids (AL, USA), was stored in chloroform at −20°C. Analytical grade chemicals were from Sigma. Silica microbeads were from Bangs Laboratories. Alexa Fluor 488 carboxylic acid succinimidyl ester dye was from Molecular Probes/Thermo Fisher Scientific. HPLC purified oligonucleotides containing the *matS19* sequence targeted by MatP (AAA**GTGACACTGTCAC**CTT, bases recognized by the protein in bold) (5), with or without fluorescein or Alexa 647 covalently attached to the 5’ end of the sense oligonucleotide, were purchased from Microsynth or IDT. Complementary strands were hybridized by heating at 85°C in a thermocycler and slowly cooling down. The fluorescently labelled oligonucleotide (*matS*-Fl or *matS*-Alexa 647) was hybridized with a 10% excess of the unlabelled complementary strand. All *in vitro* experiments were done in 50 mM Tris-HCl, 300 mM KCl, 5 mM MgCl_2_, pH 7.5 (working buffer).

### Bacterial strains and growth conditions

MG1655 *matP-mCh::kan*, a kind gift of Pauline Dupaigne (17), was grown to steady state in minimal glucose medium (Gb4: 6.33 g K_2_HPO_4_ (Merck), 2.95 g KH_2_PO_4_ (Riedel de Haen), 1.05 g (NH_4_)_2_SO_4_ (Sigma), 0.10 g MgSO_4_·7H_2_O (Roth), 0.28 mg FeSO_4_·7H_2_O (Sigma), 7.1 mg Ca(NO_3_)_2_·4H_2_O (Sigma), 4 mg thiamine (Sigma), 50 mg lysine (Sigma), 50 mg arginine (Sigma), 50 mg glutamine (Sigma), 2 mg thymidine (Sigma), 20 mg·L^−1^ Uracil (Sigma) and 4 g glucose per litre, pH 7.0) at 28°C while shaking at 205 rpm. At an OD_450nm_ of 0.2 (Biochrom Libra S70 spectrophotometer, Harvard Biosciences) cells were fixed by 2.8% formaldehyde and 0.04% glutaraldehyde for 15 min before being washed in PBS (36). After splitting in two batches, the one batch was immunolabelled with antibodies against ZapB and the other with antibodies against FtsN (16) as described (36). The nucleoids were then stained with 1 μg/mL DAPI. Secondary antibodies were donkey anti rabbit IgG conjugated to Oregon Green (Jackson Immunoresearch). When cells were imaged live, they were concentrated and resuspended gently in their own medium. Expression of pXL28 mNG-GlpT was induced for 2 mass doublings with 15 or 30 μM IPTG (Isopropyl β-D-1-thiogalactopyranoside, Duchefa) for widefield fluorescence microscopy and structured Illumination Microscopy, respectively.

### Microscopy and image analysis

For imaging the cells were immobilized on 1% agarose in water slabs on object glasses (37) and phase contrast and fluorescence microscopy images were obtained using a Nikon Eclipse Ti microscope equipped with a C11440-22CU Hamamatsu ORCA camera, an Intensilight HG 130W lamp and the NIS elements software (version 4.20.01). First a phase contrast image was taken through a CFI Plan Apochromat DM 100× oil objective, followed by a MatP-mCherry image using custom mCherry filter ex570/20, dic600LP, em605LP, a ZapB or ZapN image using GFP filter ex480/40, dic505, em535/50 and finally a DAPI image using filter ex360/40, dic400, em460/25. Images were analysed with Coli-Inspector supported by the ObjectJ plugin for ImageJ (version 1.49v) (16). Briefly, the length and diameter of more than 1200 individual cells were marked and analysed in the phase contrast images. Fluorescence and phase contrast images were aligned and fluorescence background was subtracted as described (16). The fluorescence of each cell was collected in a one-pixel wide bar with the length of the cell. A map of the diameter or the fluorescence localization and intensity was generated with the cells sorted according to increasing cell from left to right. Because cells were grown to steady state, the length of the cells can be directly correlated to the cell division cycle age. An age profile is created from all cell profiles in a map of a particular age range. They are first resampled to a normalized cell length of 100 data points, then averaged to a single plot using the macro Coli-Inspector-03s in ObjectJ (16). Concentration of the number of MatP molecules per cell and the number of molecules MatP at midcell were determined as described (16). Calculation of the Pearson coefficient of the colocalization of MatP and GlpT was determined as described (21).

### SIM sample preparation and imaging

Micron holes (1.1–1.4 μm) (38) were made with a micropillar mold in a 3% agarose in Gb4 medium layer to orient the cells vertically. Cells were concentrated by centrifugation and applied to the agarose alive. Part of the cells would enter the holes while another part would lay on top of the layer. To immobilize the cells in these holes, as MG1655 bacteria are able to rotate and move when imaged alive as they have flagella, a thin layer of 1% low melting point agarose in Gb4 medium was applied on top. A cover glass was then applied and taped to the glass slide.

SIM images were obtained with a Nikon Ti Eclipse microscope and captured using a Hamamatsu Orca-Flash 4.0 LT camera. Images were obtained with a SR APO TIRF 100x/1.49 oil objective, using 3D-SIM illumination with a 488 nm laser and an exposure time of 0.3 sec for the mNeonGreen-GlpT and a 561 nm laser with an exposure time of 1 sec for the MatP-mCherry, and were reconstructed (note that each reconstructed SIM image consists of 15 images) with Nikon-SIM software using for each picture adapted values for the parameters Illumination Modulation Contrast (IMC), High Resolution Noise suppression (HNS) and Out of focus Blur Suppression (OBS).

### MatP expression, purification and labelling

Recombinant untagged MatP was produced as previously described (17), with some modifications, from the plasmid kindly provided by Dr. M Schumacher. Briefly, the N-terminal hexa-histidine (His) tagged protein was overproduced and purified by affinity chromatography using a His-bind Resin (Novagen) with nickel. The His-tag was subsequently removed by cleavage with thrombin, followed by an ionic exchange chromatography step using a HiTrap SP HP column (GE Healthcare). The fractions of purified MatP were pooled, dialyzed against 50 mM Tris-HCl, 300 mM KCl, 1 mM EDTA, 10% glycerol, pH 7.5 and stored at −80°C. The protein concentration was measured by UV-absorbance spectroscopy using a molar absorption coefficient at 280 nm of 27960 M^−1^cm^−1^, estimated from its sequence. MatP was covalently labelled in the amino groups with Alexa Fluor 488 carboxylic acid succinimidyl ester dye (MatP-Alexa488) and stored at −80°C. The ratio of labelling was around 0.5 moles of fluorophore per mole of protein, as estimated from their molar absorption coefficients.

### Microfluidic encapsulation in microdroplets, generation of giant unilamellar vesicles and visualization by confocal fluorescence microscopy

Microfluidic devices were constructed by conventional soft lithographic techniques from masters (chip design and procedure detailed elsewhere (39)). Encapsulation was conducted at room temperature by mixing two streams of dispersed aqueous phases in a 1:1 ratio prior to the droplet formation junction. MatP (7 μM) with a tracer amount labelled with Alexa 488 (2 μM) in working buffer was one of the aqueous phases, the other one being buffer including, when stated, *matS*-Alexa 647 (2.8 μM). When present, both aqueous streams contained crowders (Ficoll or dextran). The third stream supplied the *E. coli* lipid mixture at 20–25 g·L^−1^ in mineral oil, prepared shortly before use by two cycles of vortex/sonication resuspension in the mineral oil of a lipid film obtained using a SpeedVac device. Encapsulation was also conducted including the preformed MatP *matS* complex in the two aqueous streams. Data presented correspond to experiments delivering solutions at 160 μL/h (oil phase) and 20 μL/h (aqueous phases) by automated syringe pumps (Cetoni GmbH) yielding uniform droplets. Droplets were collected during 30 min for their subsequent conversion into giant unilamellar vesicles as described elsewhere (23), introducing the outlet tubing from the microfluidic chip into 700 μL of oil phase stabilized for 1 hour over 400 μL of outer solution. Composition of the outer solution matched the encapsulated solutions, supplemented with sucrose to achieve ~25 mOsmol/Kg higher osmolarity as measured in an Osmomat 3000 (Gonotec GmbH). The solutions were then centrifuged (10–15 min, 1500 rpm in a bench centrifuge), the oil phase removed and the vesicles washed with outer solution and centrifuged again (10–15 min, 2000 rpm).

Microfluidic production of droplets on chip was monitored with an Axiovert 135 fluorescence microscope (Zeiss). The resulting microdroplets and GUVs were visualized immediately after generation by confocal microscopy with a Leica TCS-SP2 or TCS-SP5 inverted confocal microscope as previously described (23, 40). Intensity profiles in the green and red channels were obtained applying the line tool of ImageJ (National Institutes of Health) through the equatorial section of the droplets/vesicles.

### Preparation of multilamellar vesicles

Chloroform solutions of EcL were dried using a SpeedVac device. Multilamellar vesicles (MLVs) were obtained by hydration of the dried lipid film in magnesium free working buffer followed by two cycles of brief vortexing and incubation at 37°C.

### Bio-layer interferometry measurements

Lipid-protein interactions were measured by bio-layer interferometry using a single channel BLItz system (ForteBio). From the EcL MLVs, small unilamellar vesicles (SUVs) were freshly prepared before the experiments by sonication (41), and diluted in hydration buffer (50 mM Tris-HCl, 150 mM KCl, pH 7.5) to a final 0.5 g·L^−1^ concentration. Lipids were then immobilized on aminopropylsilane biosensor tips. MatP binding to the immobilized lipids was measured at the specified final protein concentrations at room temperature and with vigorous shaking (2200 rpm). Measurements were also performed in the presence of 1 μM *matS* or at variable concentration of *matS* keeping MatP at 150 nM. Assays were performed by duplicate, and binding isotherms were constructed by representing the experimental binding values at equilibrium *vs* MatP total concentration.

### Binding assays in lipid coated microbeads

Microbeads coating and binding measurements were done basically as described (42). Briefly, silica microbeads were washed, resuspended in working buffer and incubated with an excess of coating material. After removal of lipids excess and further processing to ensure even coating, microbeads were resuspended in working buffer to get the required stock concentration. The amount of lipid coating the microbeads was estimated, assuming a single bilayer and the reported value for phosphatidylcholine (43), from the surfaces ratio (gram of beads/polar head of a lipid molecule) (42). MatP binding experiments were done at constant 35 g·L^−1^ beads (62 μM accessible lipids) and variable concentration of MatP-Alexa 488. Experiments in the presence of *matS* were performed by adding, prior or after incubation with the lipids, unlabelled *matS* (0.1 or 1 μM) to the samples containing MatP-Alexa 488 (0.250 μM). Additionally, *matS*-Fl (0.1 μM) was added to samples containing unlabelled MatP (0.250 μM) and lipids. After incubation of the protein or the nucleoprotein complex with the coated beads for 20 minutes, bound material was separated by centrifugation from the free protein/nucleoprotein complex, that remained in the supernatant and was quantified using a fluorescence plate reader (Varioskan Flash, Thermo or Polar Star Galaxy, BMG Labtech) as described (42). Assays were performed by triplicate, and the binding isotherm was constructed by plotting the concentration of bound MatP as a function of concentration of free MatP. The linearity of the signal of the labelled protein with its concentration was verified.

### Analysis of protein-lipid binding isotherms

Binding parameters were obtained independently from the analysis of the binding isotherms from interferometry or fluorescence measurements by a nonlinear least-squares fit of a Langmuir adsorption isotherm:

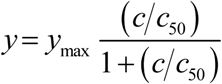

where *y* and *y*_max_ are the response and maximum response measured upon binding, respectively, *c* is the concentration of MatP and *c*_50_ is the concentration of MatP at which binding is half of the maximum value.

The method of parameter scanning (44) was employed to determine the extent to which the value of the best-fit parameter is determined by the data, as explained elsewhere (42).

## ACKNOWLEDGEMENTS

We thank M. Schumacher (Duke University) for kindly providing the MatP plasmid, X. Liu for the gift of pXL28 mNG-(GGS)_2_-GlpT, P. Dupaigne for the gift of the strain MG1655 *matP::mCh*, W.T.S. Huck and A. Piruska (Radboud University) for the kind gift of silicon masters with the chip designs, M.T. Seisdedos and G. Elvira (Confocal Laser and Multidimensional Microscopy Facility, CIB-CSIC), the Technical Support Facility (CIB-CSIC), J.R. Luque-Ortega (Analytical Ultracentrifugation and Light Scattering Facility, CIB-CSIC), N. Ropero for technical assistance, and Norbert O. E. Vischer for fruitful discussion on the analysis of foci positions.

This work was supported by the Spanish government through grants BFU2014-52070-C2-2-P and BFU2016-75471-C2-1-P (to G. R.) and by the Dutch government through grant NWO ALW open program nr. 822.02.019 (to N. Y. M.). The funders had no role in the design; data collection, analysis or interpretation; manuscript writing or the decision to submit the article for publication. The authors declare no conflicting interests.

## Author contributions

B.M., S.Z. and G.R. conceived the experimental work; B.M., S.Z., C.A., N.Y.M. and T.d.B. analysed results; B.M., S.Z., M.S.-S., M.R.-R, C.A., N.Y.M, and J.V. performed experimental work; B.M., S.Z., C.A., N.Y.M, T.d.B and G.R. discussed the results and wrote the manuscript; B.S. provided the mould to make SIM agar holes. All authors read and approved the final manuscript.

